# Characterization of Cervical-Cranial Muscle Network in Correlation with Vocal Features

**DOI:** 10.1101/2021.09.23.461177

**Authors:** Rory O’Keeffe, Seyed Yahya Shirazi, Sarmad Mehrdad, Tyler Crosby, Aaron M. Johnson, S. Farokh Atashzar

## Abstract

Objective evaluation of physiological responses using non-invasive methods has attracted great interest regarding the assessment of vocal performance and disorders. This paper, for the first time, demonstrates that the topographical features of the cervical-cranial intermuscular coherence network generated using surface electromyography (sEMG) have a strong potential for detecting subtle changes in vocal performance. For this purpose, in this paper, 12 sEMG signals were collected from six cervical and cranial muscles bilaterally. Data were collected from four subjects without a history of a voice disorder performing a series of vocal tasks. The vocal tasks were varied phonation (an /a/ sustained for the maximal duration with combinations of two levels of loudness and two levels of pitch), a pitch glide from low to high, singing a familiar song, spontaneous speech, and reading with different loudness levels. The varied phonation tasks showed the median degree, and weighted clustering coefficient of the coherence-based intermuscular network ascends monotonically, with a high effect size (|*r_rb_*| = 0.52). The set of tasks, including pitch glide, singing, and speech, was significantly distinguishable using the network features as both degree and weighted clustering coefficient had a very high effect size (|*r_rb_*| > 0.83) across these tasks. Also, pitch glide has the highest degree and weighted clustering coefficient among all tasks (degree > 0.6, weighted clustering coefficient > 0.6). Spectrotemporal features performed far less effective than the proposed functional muscle network metrics to differentiate the vocal tasks. The highest effect size for spectrotemporal features was only |*r_rb_*| = 0.19. In this paper, for the first time, the power of a cervical-cranial muscle network has been demonstrated as a neurophysiological window to vocal performance. The results also shed light on the tasks with the highest network involvement, which may be potentially used in monitoring vocal disorders and tracking rehabilitation progress.

## I. Introduction

MORE than 17 million people in the United States are estimated to suffer from dysphonia (a voice disorder) each year [1], [2]. Excessive voice use and maladaptive compensatory muscle tension in response to underlying neurological or physiological laryngeal disease are considered to be potential roots for voice disorders such as muscle tension dysphonia (MTD) [3]. The laryngeal muscles are internal to the neck and require invasive access for direct examination. However, it is suggested that the perilaryngeal cervical and cranial muscles may show activity alterations in subjects with dysphonia [4], which can help with the diagnosis using surface electromyography (sEMG).

The current methods for monitoring the function of laryngeal muscles include intramuscular EMG, external laryngeal palpation, and laryngeal endoscopy [5], [6], [7], [8], [9], [10], [11]. Although these methods have provided much information about muscle activation and function during voicing, they are invasive, uncomfortable, and subjective. Intramuscular EMG requires inserting small wires into the muscle using a needle to measure relative neuromuscular activity. Laryngeal endoscopy involves placing a flexible endoscope through the nose or a rigid endoscope through the mouth to visualize the gross anatomy and movements of the vocal folds. It requires a trained specialist to perform and subjectively interpret the findings. Manual palpation of the larynx and perilaryngeal musculature is easy to perform but does not provide any quantitative or standardized measure of muscle tension. Additionally, these evaluation methods can disturb the normal function of the muscle during the examination (for example, due to the pain), which can affect the accurate assessment.

Recording sEMG is a non-invasive technique that can potentially provide objective information about perilaryngeal muscle activity during voicing based on temporal and spectral characteristics of the muscle signal measured at the skin surface. Previous sEMG studies of the external muscles have shown inconsistent outcomes using spectrotemporal features, such as root mean square (RMS) and power spectral density (PSD). In this regard, although a recent study suggested some evidence about differences in RMS values of cervical sEMG activations between patients with muscle tension dysphonia and the control group [9], another comprehensive study did not find such differences for different types of dysphonia, including MTD [12].

In neuroscience literature, coherence analysis has been used in the context of brain connectivity to detect how different regions of the brain are synchronized (or functionally coupled) during different tasks, and this measure has been used for detecting the degrees of several central nervous system conditions, such as Parkinson’s Disease [13], [14]. More recently, using coherence analysis, the fluency of corticomuscular connectivity has also been investigated to understand how the central nervous system communicates with the peripheral nervous system [15], [16]. Similarly, the functional muscle network is an emerging concept that uses simultaneous multichannel sEMG to decode how various muscle groups are synergistically synchronized during various motor tasks [17], [18]. Intermuscular coherence networks have been recently used to holistically investigate the muscular system during various gait tasks and uniquely discriminate subtle differences in lower limb functions in non-disabled adults [19], [20].

To the best knowledge of the authors, the concept of functional muscle networks has not been used at the cervical and cranial levels. Some efforts have been conducted to assess beta-band (15-35 Hz) coherence between two anterior neck muscles during voicing, which showed some discriminative power to indicate hyperfunction and differences between control subjects and patients with vocal nodules [21], [22]. Expanding from a single coherence measurement in specific frequency bands to a wideband intermuscular coherence network increases the possibility for monitoring motor functions or impairments due to the wider spectral and spatial distribution of the analysis. Thus, it is imperative to understand the power of the cervical-cranial muscle network and the corresponding relationship with various vocal functions.

The purpose of this study is to quantify the cervical-cranial muscle network characteristics of a series of vocal tasks for healthy subjects. We hypothesize that in non-disabled subjects increasing the loudness and pitch (i.e., vocal frequency) will change the network connectivity in a manner that can be registered using topographical characteristics of the network, such as degree and clustering coefficient. In this study, to conduct a comparative analysis, the classical spectrotemporal features are also quantified to determine if the tasks with stronger muscle networks also consistently elicit statistically distinguishable spectrotemporal muscle activity. We show that the muscle network provides robust and statistically consistent discrimination for increasing loudness and pitch, suggesting that the cervical-cranial muscle network can indeed be used to detect subtle differences in vocal tasks, while the conventional spectrotemporal features fail to function accordingly.

## II. Methods

Four healthy subjects (all males, 39.5 ± 7 years) participated in the study. The institutional review board of the New York University Grossman School of Medicine approved the study, and subjects provided their written consent after they received the study description. Subjects denied any history of dysphonia or neck and cervical-related injuries.

### A. Experimental Procedure

Subjects performed a series of vocal tasks, each of a different type or while varying tonal parameters (Fig. 1b). The first group of tasks involved making a maximally sustained /a/ sound at a constant pitch and volume. With two levels of loudness and two levels of pitch, in total, there were four varied phonation (/a/ sound) tasks: 1) habitual loudness, habitual pitch, 2) elevated loudness, habitual pitch, 3) habitual loudness, high pitch and 4) elevated loudness, high pitch. Subjects were instructed to sustain the /a/ sound for as long as was comfortable. Subjects performed three trials of each of loudness and pitch combinations before moving to the next tasks. The second group of tasks included single repetition vocal exercises, namely (i) pitch glide, (ii) spontaneous speech, and (iii) singing. Pitch glide involved starting to intone at a low pitch and smoothly increasing to a final high pitch [23]. The spontaneous speech task involved responding to the prompt, “tell me how to make a peanut butter and jelly sandwich,” in a typical conversational voice, while the singing task involved singing ‘Happy Birthday’ in a comfortable key chosen by the participant. The third group of tasks involved reading the first full paragraph of The Rainbow Passage [24], a standard reading passage used to evaluate the voice, at three levels of loudness: habitual, elevated, and whispering.

**Fig. 1.**
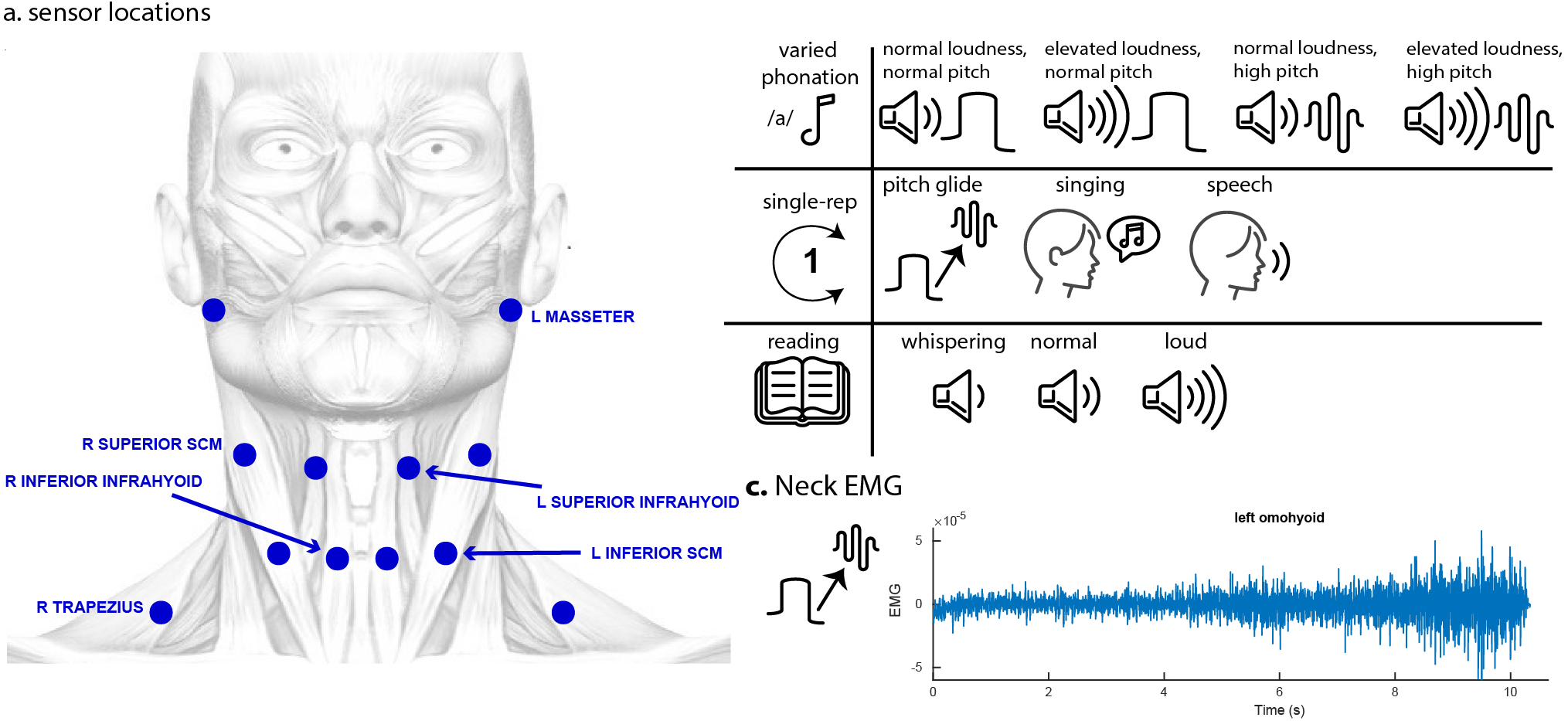
**a.** Six sensors were placed on each side of the neck at Masseter, Superior Sternoclediomastoid (Superior SCM), Superior Infrahyoid, Inferior Infrahyoid, Inferior Sternocleidomastoid (Inferior SCM) and Trapezius. **b.** Subjects performed vocal tasks, classified as varied phonation, single repetition, or reading. Varied phonation tasks involved intoning /a/ at defined loudness and pitch levels. Single repetition tasks included pitch glide, singing, and speech. Reading tasks involved reading a passage at different loudness levels. **c.** An example of a neck sEMG signal trace, taken from left superior infrahyoid for Subject 4 during pitch glide. Note how the amplitude level increases as the task develop.

sEMG signals were recorded from twelve sensors, using the wireless Trigno sEMG system (Delsys Inc., Natick, MA), with a sampling frequency of 1259 Hz (Fig. 1a). Four bipolar Trigno Mini sensors were used for the inner cervical muscles (inferior and superior infrahyoid, bilaterally), while eight bipolar Trigno Avanti sensors were used for Masseter, Superior Sternocleidomastoid, Inferior Sternocleidomastoid, and Trapezius. With regard to palpation, subjects were instructed to (i) clench their teeth to identify masseter, (ii) look left and right to identify lower and upper sternocleidomastoid, (iii) look up and down to identify the infra-hyoid muscles, (iv) move shoulders forwards and backward before staying a neutral position to identify trapezius muscles. The skin surface was thoroughly wiped prior to sensor placement. Sensors were placed parallel to the direction of the muscles. In order to minimize the noise content of the recorded signals, subjects were instructed not to move their head during the task. Following the recording, signals were pre-processed using MATLAB R2020b (MathWorks Inc. Natick MA). The first and last 1s of all trials were clipped out, and other trials were clipped further in the case of a head movement at the beginning or at the end. Afterward, a zero-phase Butterworth high pass filter at 1 Hz, a zero-phase Butterworth bandstop filter between 57.5-62.5 Hz, and a zero-phase Butterworth low pass filter at 110 Hz were all applied. An example of a cervical-cranial muscle signal (from left superior infrahyoid during pitch glide) is shown in Fig. 1c.

### B. Muscle Signal Analysis

Muscle networks were constructed for all tasks, using coherence. Magnitude squared coherence, *C_xy_* between two signals *x*(*t*) and *y*(*t*) is:

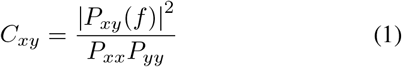

where *P_xx_* and *P_yy_* are the power spectral densities (PSDs) and *P_xy_* is the cross power spectral density (CPSD). To compute the coherence, Welch’s overlapped averaged periodogram method [25] was utilized with a Hamming window of 2048 samples (1.63 ms) and 50% overlap. The maximum coherence component in the 5-100 Hz range was selected for each sensor pair. Using this maximum coherence value, muscle networks were constructed for each trial. Each node in the network represents a muscle, and the width of each line illustrates the pairwise muscle coherence. In the case of tasks that had multiple trials, the median network across trials was computed. The degree of each node, *D_i_*, is the average of all edges connected to the node. If the muscle network is represented by adjacency matrix *A, D_i_* is defined as:

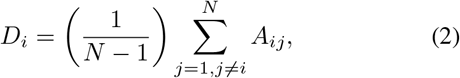

where *N* is the number of nodes. A node’s weighted clustering coefficient (*WCC_i_*) gives the measure of how well that node is connected to its neighbors. The weighted clustering coefficient is defined as:

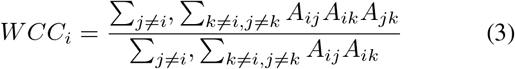

A node that is not connected to its neighbors will have a weighted clustering coefficient, *WCC_i_* = 0, while a node that is very well connected to its neighbors has *WCC_i_* = 1.

In order to provide a comparison between the muscle coherence network and the conventional spectrotemporal metrics, muscle activations were quantified in the time and frequency domains. The time-domain activation was quantified by finding the RMS value across the trial duration. With regard to the spectral domain, PSD was computed using Welch’s method [25] and the median PSD across 5-100 Hz was computed. Furthermore, the median frequency was computed for each task. The median frequency is defined as the frequency at which the area under the PSD graph is divided in two. The median value across trials was used for PSD, median frequency, and RMS when there were multiple trials.

### C. Statistical Analysis

In order to evaluate the statistical trends observed in coherence muscle networks, a coherence distribution was constructed for each task (Fig. 4). Each distribution consisted of degree and weighted clustering coefficient for all nodes across all subjects’ muscle networks, giving *n = #subjects × #nodes* = 4 × 12 = 48. Similarly, distributions were constructed for RMS, PSD, and median frequency (*n* = 48 for all). The Kolmogorov-Smirnov test for normality rejected the normal distribution hypothesis for the coherence, RMS, PSD, and median frequency distributions. Therefore, nonparametric statistical tests were used in our analysis. The Friedman test was used to compare tasks in each group (i.e., varied phonation, single-repetition, reading). The Wilcoxon signed-rank test was used as a posthoc test if the Friedman test revealed significance. The significance level, *α*, for all tests was initially set at 0.05. To adjust for multiple comparisons, the Bonferroni correction was applied, dividing *α* by the number of comparisons.

Finally, by using the rank-biserial correlation, the effect size of the non-normal distributions was quantified [26]. In this regard, |*r_rb_*| was used for measuring the rank-biserial correlation. A higher value means that the effect size is larger. For example, as can be seen in Fig. 4b, the coherence degree for single repetition tasks has a very high effect size (|*r_rb_*| = 0.85), and the difference between tasks is even visually clear. On the other hand, the coherence degree for reading tasks has a low effect size (|*r_rb_*| = 0.08), as there is not a clear relationship between coherence and reading task loudness.

## III. Results

### A. Coherence Networks

The median network across subjects displays a visible difference between vocal tasks, e.g., between pitch glide and speech (Fig. 2, middle row left vs. middle row right). Similarly, Fig. 3 shows that the mean degree of the network changed by the task for all subjects. Interestingly, the mean degree showed a monotonic increasing trend in response to both raised loudness and pitch for the varied phonation tasks, and pitch glide appears to have the highest coherence of all 10 tasks (Fig. 3). Mean degree showed a monotonically decreasing trend from pitch glide to singing to speech. There appears to have been little to no difference in mean degree observed for reading tasks with different loudnesses (Fig. 3).

**Fig. 2.**
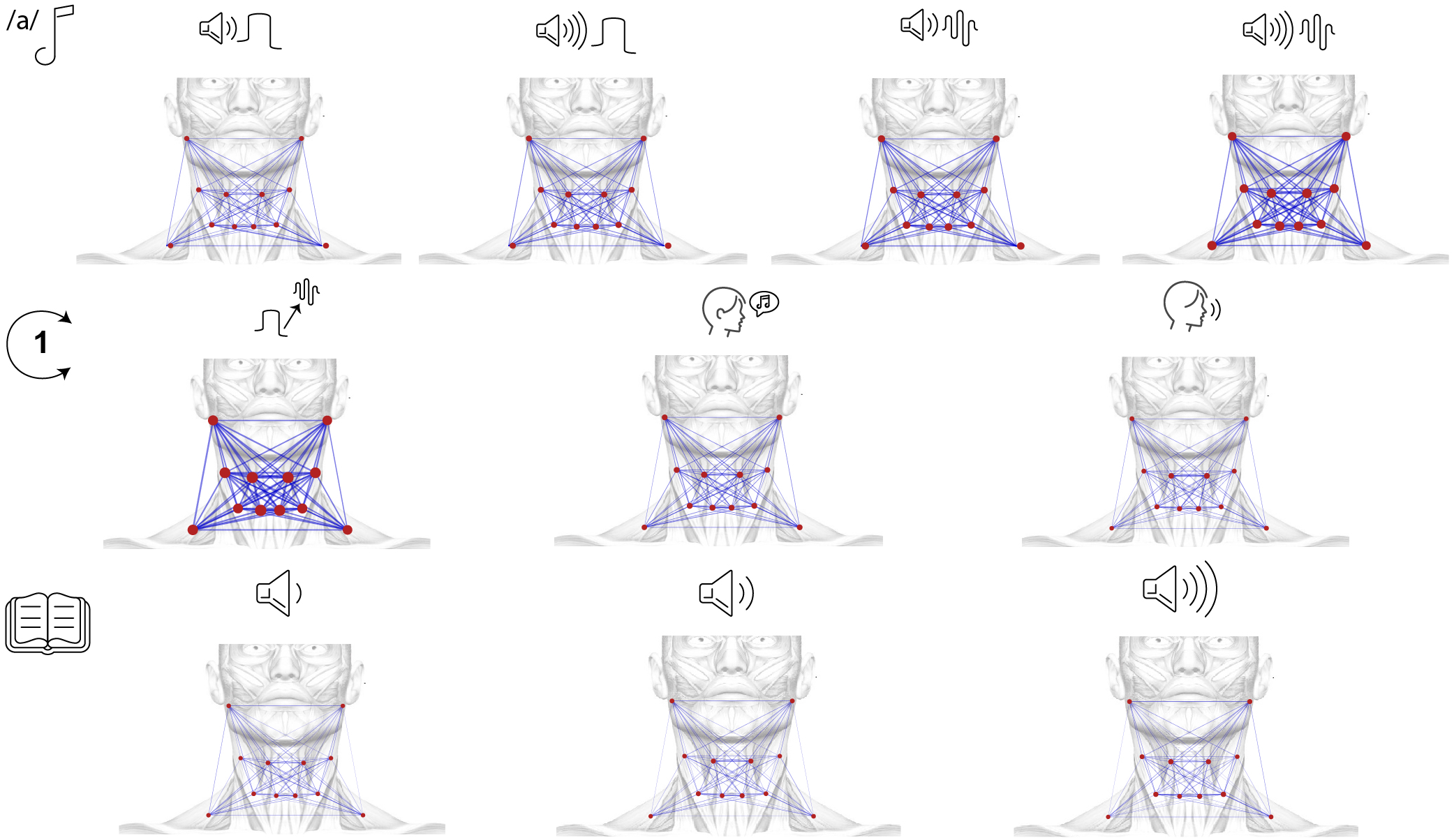
Coherence muscle networks for each vocal task were created from twelve sEMG sensors placed bilaterally on the neck area. The width of each line denotes the pairwise coherence between the two connected muscles, which is equal to the maximum coherence component in the 5-100 Hz range. In the case of tasks that had multiple trials, the median network across trials is shown. Each node radius is equal to the degree (mean of coherences involving that node). The line widths and node radii seem largest for /a/ with elevated loudness, high pitch, and pitch glide tasks.

**Fig. 3.**
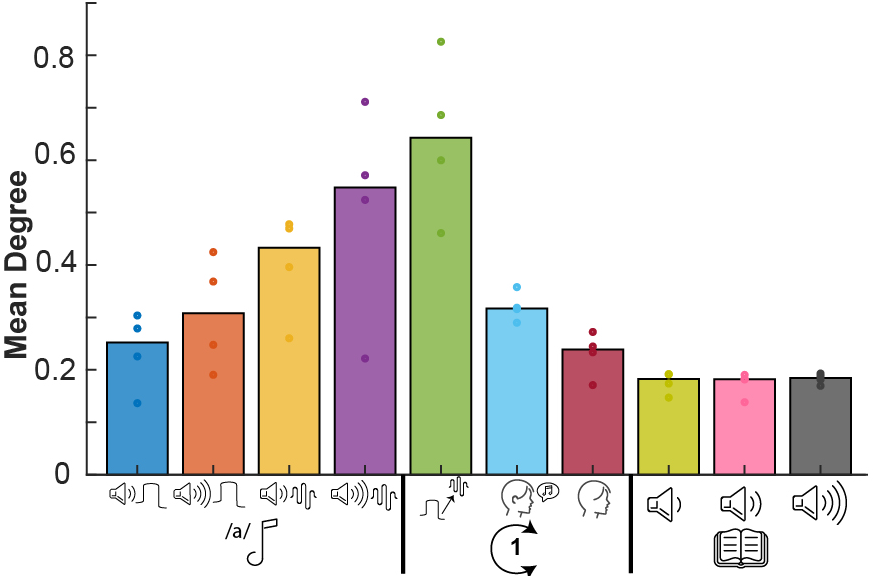
network mean degree for each task. The bar depicts the median value across subjects. Individual subject values are denoted by dots. For the subject median, the mean degree shows an increasing trend starting from the first task (habitual loudness, habitual pitch) and continuing incrementally until pitch glide. Network connectivity then shows a decreasing trend from pitch glide to singing to speech. Finally, reading tasks seem to have little difference between each other.

In order to support the initial observations of the coherence network differences between the vocal tasks, coherence distributions were constructed by including all nodes in the subjects’ intermuscular network, measured using degree and weighted clustering coefficient (Fig. 4). For the varied phonation tasks, the task-wise network degree and weighted clustering coefficient median were monotonically ascending with increasing pitch and loudness, and **all** tasks were statistically different from each other (*Friedman* 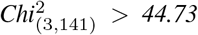, *p < 0.05, posthoc Wilcoxon signed-rank test: **all six** pairwise comparisons p* < 0.001). The network’s global efficiency showed a trend of monotonically increasing coherence with pitch and loudness for 3 out of 4 subjects. Moreover, the degree’s effect size indicated a high value of |*r_rb_*| = 0.52. For the single repetition tasks, the median of network degree and weighted clustering coefficient was decreasing monotonically, with the pitch-glide having the highest network degree and weighted clustering coefficient at both greater than 0.6 (*Friedman* 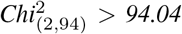, *p < 0.05, post-hoc Wilcoxon signed-rank test for **all three** pairwise comparisons, p < 0.001*).

**Fig. 4.**
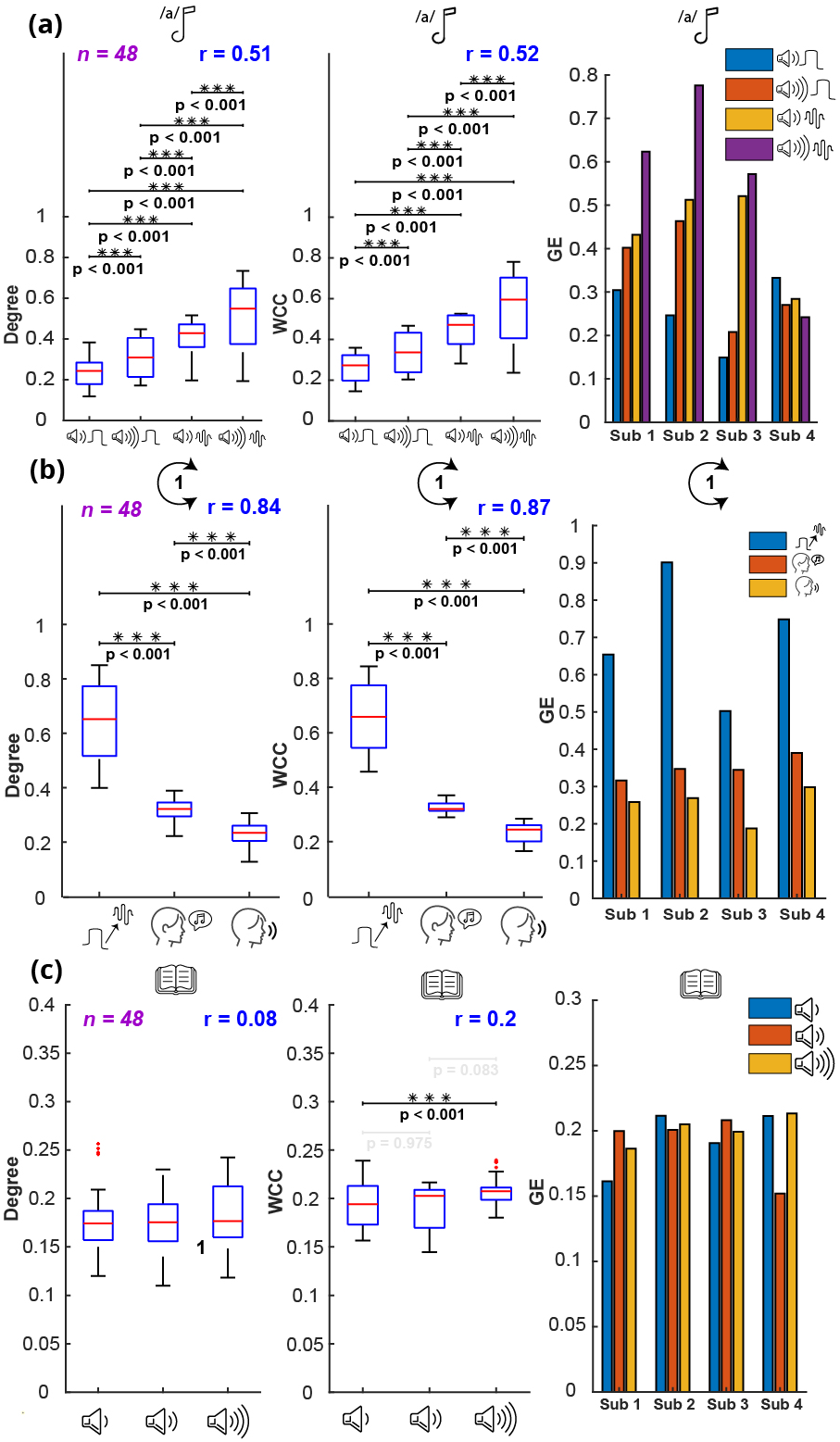
Statistical results for network metrics of the three groups of tasks. The coherence muscle network was constructed for each task, and produces 12 node values for each of degree and weighted clustering coefficient (WCC). For each distribution of degree and weighted clustering coefficient, all node values for all subjects are included. For global efficiency, a singular value was obtained for each subject’s muscle network. The columns are organized as follows, from left to right: (i) the degree distribution (*n* = 48), (ii) weighted clustering coefficient distribution (*n* = 48) and (iii) bar plots comparing subjects’ trends for taskwise global efficiency. **(a)** For **varied phonation** tasks, intermuscular network degree and weighted clustering coefficient increase monotonically with raised loudness and pitch, with statistical significance (all *six taskwise comparisons:* p < 0.001). The effect size (|*r_rb_*|) is quite high for both degree (|*r_rb_*| = 0.51) and weighted clustering coefficient (|*r_rb_*| = 0.52). **(b)** For **single repetition** tasks, intermuscular network degree and weighted clustering coefficient decrease monotonically from pitch glide, to singing, to speech, with statistical signficance (*all three taskwise comparisons:* p < 0.001). The effect size (|*r_rb_*|) is very high for both degree (|*r_rb_*| = 0.84) and weighted clustering coefficient (|*r_rb_*| = 0.87). All subjects’ global efficiency bar plots follow the monotonically decreasing pattern from pitch glide to singing to speech. **(c)** For **reading** tasks, there are no visible trends in degree, weighted clustering coefficient or subject-wise global efficiency. There are no statistically significant differences between network degree of reading tasks. For weighted clustering coefficient, there is a significant difference (*p* < 0.001) between whispering and loud reading. Effect size for degree (|*r_rb_*| = 0.08) and weighted clustering coefficient (|*r_rb_*| = 0.2) is low.

Furthermore, the global efficiency trend was consistent and decreasing across all four subjects. The degree’s rank biserial correlation also indicated a very high effect size (|*r_rb_*| =0.84). For the reading tasks, the Friedman test did not show a significant difference for the network degree but was significant for the weighted clustering coefficient (network degree: *Friedman* 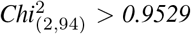, weighted clustering coefficient: *Friedman* 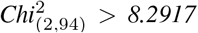. The post-hoc Wilcoxon signed-rank test indicated that only the whispering and loud reading were significantly different (*p* < 0.001). Global efficiency did not indicate a consistent trend amongst subjects for the reading tasks. Moreover, for the reading task set, the effect size was low for the network degree (|*r_rb_*| = 0.08) and weighted clustering coefficient |*r_rb_*| = 0.2

Since one of the secondary aims of this study is to suggest a suitable candidate task(s) for monitoring the effect of therapy, the tasks which had produced the highest network metrics were identified. For this, three tasks were selected, which had resulted in the highest response in Fig. 4, when compared within their categories. The selected three tasks are (i) the varied phonation task with elevated loudness, and high pitch, (ii) pitch glide, and (iii) loud reading, and results are given in Fig. 5. Both network degree and weighted clustering coefficient were significantly different from each other for **all** comparisons (*Friedman* 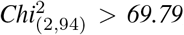, *p < 0.05, post-hoc Wilcoxon signed-rank test: **all three** pairwise comparisons p < 0.001*). Moreover, pitch glide had the highest network degree (median value ~ 0.65) and weighted clustering coefficient (median value ~ 0.7), even higher than the varied phonation task with elevated loudness and high pitch (degree: median value ~ 0.55, weighted clustering coefficient: ~ 0.6). Reading at elevated loudness had a lower degree (median value ~ 0.17) and weighted clustering coefficient (median value ~ 0.19) than the other two tasks. This suggests that pitch glide generates the maximum response of the network, which can be considered potentially the most responsive and suitable task for identifying abnormalities.

**Fig. 5.**
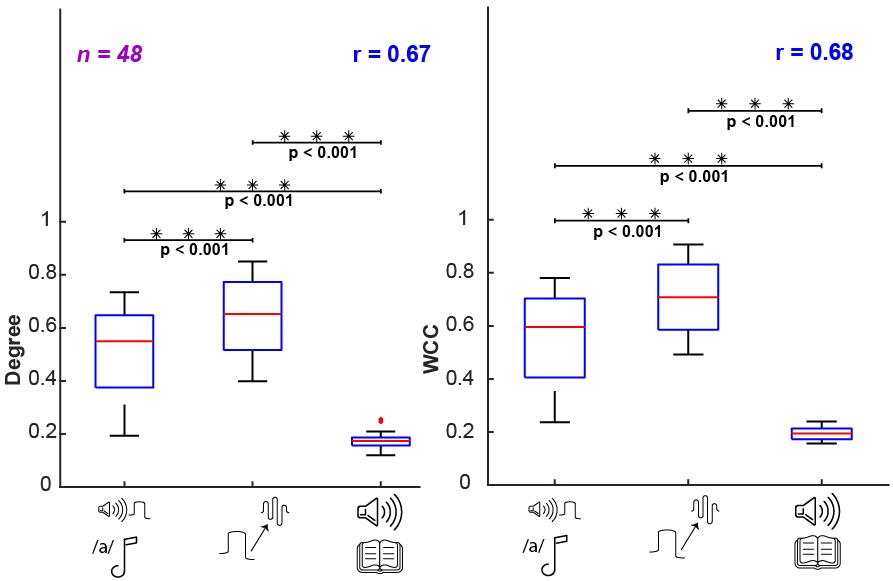
Comparison of tasks with highest network metrics from each group: 1) Pitch glide has both the highest degree and weighted clustering coefficient (WCC), followed by 2) varied phonation task with elevated loudness, high pitch, followed by 3) loud reading, and all tasks are different from one another (*for both degree and weighted clustering coefficient, all three taskwise comparisons: p* < 0.001). Reading has ~ 1/3 the median degree and weighted clustering coefficient of the varied phonation task and pitch glide.

### B. Spectrotemporal Metrics

To compare the ability of spectrotemporal metrics to distinguish different tasks, statistical analyses on RMS, PSD, and median frequency of sEMG were conducted. Distributions for each of the three aforementioned quantities were constructed by considering all nodes across all subjects (*n=#subjects x #nodes = 48*). For the varied phonation tasks, the median RMS of louder tasks is higher than habitual loudness tasks (*p* < 0.001). In this regard, PSD shows the same trend as RMS (*p* < 0.036). However, the overall effect size of varied phonation tasks (RMS: |*r_rb_*| = 0.13, PSD: |*r_rb_*| = 0.12, median frequency: |*r_rb_*| = 0.13) is quite small. For single repetition tasks, RMS and PSD did not show a clear trend (Fig. 6b). However, median frequency of pitch glide was higher than other tasks (*p* < 0.001), |*r_rb_*| = 0.16. For reading tasks, PSD and RMS are weakly correlated with loudness (Fig. 6c). All distributions are different from each other (RMS: *p* < 0.026, PSD: *p* < 0.006) and the rank biserial correlation, |*r_rb_*| = 0.19, suggests a weak effect size.

**Fig. 6.**
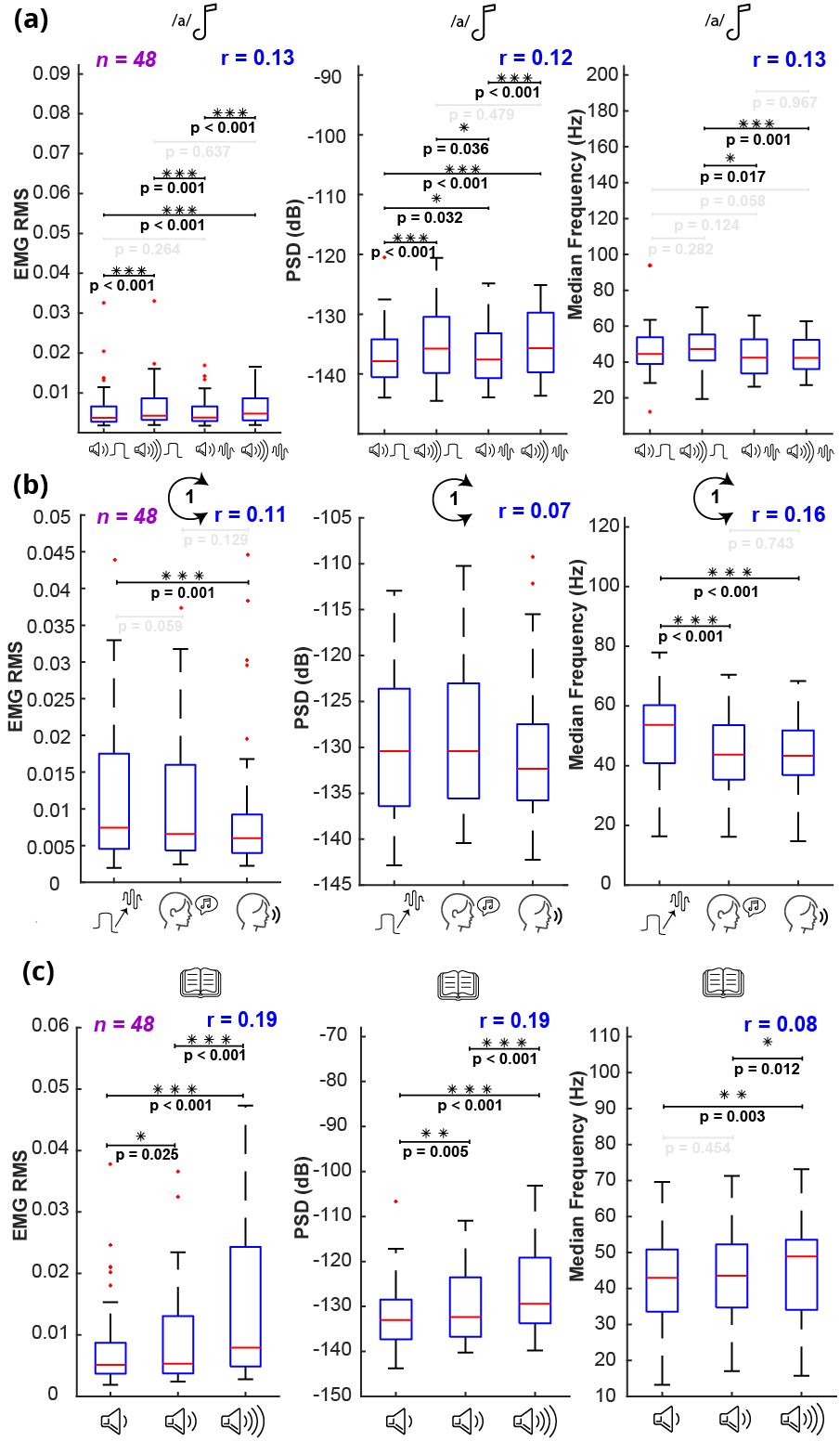
Statistical results for **RMS, PSD** and **median frequency** of the three groups of tasks. Quantifying the muscle activity with RMS, PSD (median across 5-100 Hz) and median frequency produces 12 node values for each metric. All node values for all subjects are included in each distribution. The columns are organized as follows, from left to right: (i) sEMG RMS distribution (*n* = 48), (ii) the power spectral density (PSD) distribution (*n* = 48), (iii) median frequency distribution (n = 48). **(a)** For **varied phonation** tasks, RMS and PSD show increases in response to raised loudness. Louder tasks have higher RMS (*p* < 0.001) and PSD (*p* < 0.036) than habitual loudness tasks. However, overall effect size of varied phonation tasks was still quite low (RMS: |*r_rb_*| = 0.13, PSD: |*r_rb_*| = 0.12). No consistent patterns were observed for median frequency. **(b)** For **single repetition** tasks, pitch glide had a significantly higher median frequency than the two other tasks (*p* < 0.001). No substantial trends were observed for RMS or PSD. **(c)** For **reading** tasks, there is an apparent pattern of increasing RMS and PSD in response to raised reading loudness. However, the effect size is weak (RMS and PSD: |*r_rb_*| = 0.19).

## IV. Discussion

The results show that the cervical-cranial intermuscular coherence network can distinguish both changes in vocal parameters (i.e., loudness and pitch) and different vocal tasks. The muscle network quantifies the spectral synchrony and can capture subtle changes associated with vocal output. This is demonstrated initially by the network visualization, and then statistical analysis using the network metrics (degree and weighted clustering coefficient) confirmed the observed trends. With regard to the vocal tasks, two important statistically robust trends are identified as follows: (i) network degree and weighted clustering coefficient increase monotonically with loudness and pitch in the varied phonation tasks and (ii) network degree and weighted clustering coefficient are both the highest for the pitch glide task. The ability of the intermuscular coherence network to distinguish vocal parameters and tasks was far superior to the conventional node-wise metrics for sEMG, such as RMS and PSD. These results suggest that both the varied phonation and single repetition tasks in combination with the cervical-cranial myographic network are sensitive to various vocal features and thus can be potentially considered as candidates for measuring the efficacy of therapy for vocal disorders. The current study showed very strong statistics supporting the use of the proposed network measures while the conventional spectrotemporal metrics fail to provide the needed sensitivity to the vocal features.

Intermuscular cervical-cranial coherence ascends monotonically with loudness and pitch in the varied phonation tasks (Figs. 2, 3 and 4a). The high differentiation of the varied phonation and single repetition group tasks with high to very high effect size (varied phonation: |*r_rb_*| = 0.52, single repetition: |*r_rb_*| = 0.84) robustly supports the hypothesis that cervical-cranial intermuscular coherence is generally proportional to loudness and pitch. To the best of the authors’ knowledge, this is the first study that shows cervical-cranial intermuscular coherence is conclusively correlated to loudness and pitch.

In contrast to the coherence network, conventional nodewise metrics such as RMS, PSD, and median frequency failed to efficiently discriminate varied phonation tasks (Fig. 6a). Although there were some differences between louder and habitual loudness tasks, the effect size (|*r_rb_*|) was much smaller for these metrics than for coherence (RMS: 0.13, PSD: 0.12, versus network degree and weighted clustering coefficient: 0.52), i.e., the average difference between the task pairs was less pronounced compared to the median of the tasks. The superior performance of coherence over nodewise metrics may arise from the fact that network analysis allows us to conduct a holistic neurophysiological analysis of the functional synchrony and synergistic co-modulation of the muscles needed for successful conduction of the tasks. Thus, in the context of voice, the authors believe that the synchronous behavior of the muscles has higher discriminative power (for separating vocal features) and potentially higher diagnostic value than isolated individual muscle recordings.

Overall, the single repetition tasks provided the most differentiable network degree and weighted clustering coefficient, which both decrease monotonically from pitch glide to singing to speech with very high effect size (degree: |*r_rb_*| = 0.82, weighted clustering coefficient: |*r_rb_*| = 0.87) (Fig. 5). The effect is even greater for the single repetition than varied phonation tasks. Moreover, it should be highlighted that all subjects followed the group trend for network global-efficiency of the single repetition tasks, emphasizing robust separability. This suggests that vocal tasks of different nature (pitch glide, singing, and speech) provide the greatest diversity and objectivity of muscle network performance which can be easily differentiated by the network metrics. The fact that pitch glide has the highest degree and weighted clustering coefficient among all tasks (Fig. 6) suggests that the smooth transition of the voice through octaves is assisted by very synchronous cervical-cranial muscle activity. Indeed, it is notable that singing had higher network coherence than regular speech. Using the results that (i) higher pitch led to increased network coherence for /a/ tasks and (ii) pitch glide has the maximum network coherence of all tasks, both the higher average pitch and the larger number of pitch changes for singing versus speech are consistent with singing having higher network coherence. This result is in contrast to a previous result with beta-band coherence between two muscles [21], which found that speech had higher beta-band coherence than singing. This difference might be due to benefiting from an intermuscular coherence network with 12 nodes and 66 edges in this study versus only two nodes and one edge coherence in the previous study. This work also considers a much wider frequency range (5-100 Hz) than the previous beta-band coherence [21]. Despite the higher expected pitch for singing vs. speech, neither median frequency nor PSD succeeded in detecting a difference here, highlighting the superiority of muscle network coherence over node-wise methods in responding to subtle physiological changes of external muscles related to vocal output.

Tasks with a higher pitch component produced a more synchronous network, and varying the pitch led to clearer network responses. Both the varied phonation (/a/) high pitch and pitch glide tasks were shown to have at least 3 × the network degree or weighted clustering coefficient of a reading task (Fig. 5), highlighting the ability of tasks with a high pitch component to produce the most pronounced network output. Moreover, the pitch-varying task sets showed clear responses to vocal parameter changes. The varied phonation tasks showed a gradually increasing response (Fig. 4a) from a median network degree ~ 0.2 for habitual loudness, habitual pitch to ~ 0.6 for elevated loudness, high pitch. The single repetition tasks showed a sharper decrease from pitch glide to singing than from singing to speech. For both varied phonation and single repetition tasks, the network response to each task was distinctive; the effect size was large (|*r_rb_*| > 0.52), and the null hypothesis was rejected with the highest significance level (*p* < 0.001) for all task-wise comparisons. Tasks that produce the most responsive network behavior would be the most appropriate candidates in vocal therapy assessment, as the responsiveness of the cervical-cranial intermuscular coherence network should be high to maximize the sensitivity of detecting signs of dysphonia or improvements made by the therapy. Since the muscle network was most responsive to tasks with a higher pitch component and pitch-varying task sets, such tasks promise the best chance of success when monitoring patient progress during vocal therapy.

Our results demonstrate the efficacy and sensitivity of the intermuscular coherence network analysis in reflecting subtle modulations in vocal output, such as detecting changes in vocal parameters and discriminating single repetition tasks, with a robustly high effect size. Taking inspiration from brain connectivity networks that can detect functional changes, this is the first work that shows topographical features of the intermuscular coherence network can detect changes to indicate functional vocal characteristics. This will be particularly beneficial to detect unhealthy differences in vocal activity caused by physiological damage. Our study showed that the high functional synchronicity of the cervical-cranial muscles produced a strong network response during pitch glide, suggesting that the laryngeal performance can be measured by the cervical-cranial network, which needs to function in synchrony to conduct the corresponding tasks. With such a high sensitivity, other vocal disorders could be potentially detected and monitored by our suggested configuration. Given the high range of muscles recorded, in addition to the wide frequency range covered, the cervical-cranial muscle network is a great candidate for an objective, digital method of detecting and monitoring a wide range of vocal disorders, using smart wearable clinical technologies, such as a smart EMG necklace. A further clinical application of the strong correlation between cervical-cranial muscle network features and vocal output lies in an EMG-based electrolarynx device [27], [28]. Regarding patients who have lost their vocal cords due to cancer, the muscle network features of healthy cervical muscle activity could be used to drive a laryngeal prosthesis, capable of distinct levels of output pitch and loudness.

Limitations of this study were that we didn’t control for the effect of vocal fatigue and the order of the tasks was not randomized; therefore, tasks later in the session could have been influenced by fatigue more than earlier ones. However, it should be noted that the vocal experiment procedure for each subject required ~ 9 minutes of vocal effort, which is significantly lower than the effort needed for fatiguing a subject (which is ~ 60 minutes for vocal fatigue with comfortable reading loudness or intermittent loud reading tasks for several hours [29], [30].

In conclusion, this work for the first time shows that the cervical-cranial intermuscular coherence network can detect subtle changes distinguishing vocal tasks. The network showed a robust effect size for changes in loudness and pitch in a set of varied phonation tasks and an even more robust effect size amongst single repetition tasks, which included a pitch glide, singing, and a short speech. The network out-performed conventional spectrotemporal node-wise metrics (RMS, PSD, and median frequency) regarding sensitivity to changes in vocal output. The responsiveness of the proposed network to either varied phonation or single repetition tasks, as well as its high range of muscles recorded, suggests that monitoring cervical-cranial intermuscular coherence shows promise as a method to also differentiate physiological abnormalities in patients with a wide range of vocal disorders, and optimize their therapeutic regimen.

